# Adenosinergic modulation of layer 6 microcircuitry in the medial prefrontal cortex is specific to presynaptic cell type

**DOI:** 10.1101/2023.12.04.569880

**Authors:** Chao Ding, Danqing Yang, Dirk Feldmeyer

**Affiliations:** Research Center Juelich, Institute of Neuroscience and Medicine 10, Research Center Juelich, 52425 Juelich, Germany; Department of Psychiatry, Psychotherapy, and Psychosomatics, RWTH Aachen University Hospital, 52074 Aachen, Germany; Jülich-Aachen Research Alliance, Translational Brain Medicine (JARA Brain), Aachen, Germany; Synaptic Transmission in Energy Homeostasis Group, Max Planck Institute for Metabolism Research, 50931 Cologne, Germany

## Abstract

Adenosinergic modulation in the PFC is recognised for its involvement in various behavioural aspects including sleep homoeostasis, decision-making, spatial working memory and anxiety. While the principal cells of layer 6 (L6) exhibit significant morphological diversity, the detailed cell-specific regulatory mechanisms of adenosine in L6 remain unexplored. Here, we quantitatively analysed the morphological and electrophysiological parameters of L6 neurons in the rat medial prefrontal cortex (mPFC) using whole-cell recordings combined with morphological reconstructions. We were able to identify two different morphological categories of excitatory neurons, those with a leading dendrite that was oriented either upright (towards the dial surface) or inverted (towards the white matter). These two excitatory neuron subtypes exhibited different electrophysiological and synaptic properties. Adenosine at a concentration of 30 µM indiscriminately suppressed connections with either an upright or an inverted presynaptic excitatory neuron. However, using lower concentrations of adenosine (10 µM) revealed that synapses originating from L6 upright neurons have a higher sensitivity to adenosine-induced inhibition of synaptic release. Adenosine receptor activation causes a reduction in the probability of presynaptic neurotransmitter release that could be abolished by specifically blocking A_1_ adenosine receptors (A_1_ARs) using 8-cyclopentyltheophylline (CPT). Our results demonstrate a differential expression level of A_1_ARs at presynaptic sites of two functionally and morphologically distinct subpopulations of L6 principal neurons, suggesting that they may play distinct roles in the maintenance of sleep homoeostasis by adenosine.

## Introduction

Adenosine is a nearly ubiquitous endogenous neuromodulator involved in sleep homoeostasis and energy metabolism of neurons (Ribeiro, Sebastiao et al. 2002, Porkka-Heiskanen and Kalinchuk 2011). During prolonged wakefulness, hypoxia, ischemia or elevated metabolic demand, the extracellular adenosine concentrations can rise from nanomolar to micromolar levels indicating its critical role in both physiological and pathological CNS function (Dunwiddie and Masino 2001, van Calker and Biber 2005, Fredholm 2007). Among the four types of adenosine receptors, the A_1_AR has the highest adenosine affinity and is the most abundant adenosine receptor type in the brain. It is densely expressed in the neocortex, particularly in layers 4 and 6 (Dunwiddie and Masino 2001, Hackett 2018). Acting via A_1_AR, adenosine not only reduces the intrinsic excitability of glutamatergic neurons by increasing inwardly rectifying potassium conductance (Rainnie, Grunze et al. 1994, Luscher, Jan et al. 1997) but also depresses excitatory neurotransmitter release by reducing presynaptic calcium influx and transmitter release (Wu and Saggau 1994, Arrigoni, Rainnie et al. 2001, Kerkhofs, Xavier et al. 2017, Qi, van Aerde et al. 2017). Therefore, the historical perspective has depicted adenosine as a primary inhibitory neuromodulator, exerting its function by suppressing neuronal activity through A_1_ARs within the CNS. However, more recent research has revealed that adenosine also plays a pivotal role in augmenting excitatory synaptic transmission via acting A_2A_ARs across many brain regions (Cunha and Ribeiro 2000, Ciruela, Casado et al. 2006, Rebola, Lujan et al. 2008, Simoes, Machado et al. 2016, Kerkhofs, Canas et al. 2018). While A_1_ARs mainly contribute in shaping short-term synaptic plasticity, A_2A_ARs are more engaged in modulating long-term potentiation (d’Alcantara, Ledent et al. 2001, de Mendoncca and Ribeiro 2001, Qi, van Aerde et al. 2017, Lopes, Goncalves et al. 2023). In light of this, adenosine is a neuromodulator that contributes to the precision of information processing mediated by a balanced activation of the inhibitory A1 and the excitatory A_2A_ARs.

The effects of adenosine on cortical neurons and networks across different brain regions have been well investigated (Lopes, Cunha et al. 1999, Arrigoni, Chamberlin et al. 2006, Yang, Franciosi et al. 2013, Bannon, Zhang et al. 2014, van Aerde and Feldmeyer 2015, Zhang, Bannon et al. 2015, Qi, van Aerde et al. 2017). Heterogeneous modulation of adenosine has been found in the hippocampus, striatum, somatosensory cortex and cortico-striatal pathways, indicating its cell type-specific regulatory effects (Yoon and Rothman 1991, Quiroz, Lujan et al. 2009, Rombo, Dias et al. 2016, Qi, van Aerde et al. 2017). Cortical layer 6 excitatory neurons are known to exhibit a high degree of morphological diversity in many neocortical areas including the mPFC (Yang, Seamans et al. 1996, Zhang and Deschênes 1997, Kumar and Ohana 2008, Thomson 2010, Pichon, Nikonenko et al. 2012, Marx and Feldmeyer 2013, van Aerde and Feldmeyer 2015, Yang, Qi et al. 2022). However, the intricate cell-specific regulatory mechanisms of adenosine within layer 6 (L6) have yet to be studied.

The role of adenosine in sleep homoeostasis is well documented. It has been shown that adenosine can promote and maintain sleep (Porkka-Heiskanen, Strecker et al. 1997, Strecker, Morairty et al. 2000, Basheer, Strecker et al. 2004). Adenosine receptor antagonists such as caffeine can promote wakefulness and influence normal sleep patterns. Among many higher-order brain regions, the prefrontal cortex (PFC) is particularly sensitive to sleep and benefits from it in important ways (Oniz, Inanc et al. 2019). Sleep deprivation leads to a loss of functional connectivity in the PFC and has a particular impact on brain functions that are mediated by the PFC, such as decreased working memory and increased irritability and apathy (Harrison, Horne et al. 2000, Verweij, Romeijn et al. 2014). In addition, it has been demonstrate that A_2A_ARs play a regulatory role in the adenosinergic modulation of PFC activity and are involved in various behavioural aspects, such as decision-making, spatial working memory and anxiety (Caetano, Pinheiro et al. 2017, Leffa, Pandolfo et al. 2018, Li, Chen et al. 2018). To gain a more comprehensive understanding of the underlying mechanisms, it is crucial to investigate the adenosinergic modulation of synaptic transmission in the PFC.

Here, we performed a quantitative classifications of excitatory neurons in L6 of rat medial PFC (mPFC) by using unsupervised cluster analysis. Based on their dendritic morphology, two main clusters of L6 excitatory neurons were identified : (1) upright excitatory neurons with their leading or apical dendrites spanning superficial layers and (2) inverted/horizontally oriented excitatory neurons with their leading dendrites projecting towards the white matter or horizontally to it. Following this, we investigated the adenosinergic modulation of synaptic connections established by these L6 excitatory neuron types using paired patch clamp recordings with simultaneous biocytin fillings. We found that connections with a presynaptic upright excitatory neurons were more sensitive to low concentrations of adenosine (10 µM) than connections with presynaptic inverted cells, although high concentrations of adenosine (30 µM) strongly suppressed all L6 excitatory connections via A_1_AR activation. Our results revealed a cell type-specific effect of adenosine on the synaptic transmission in L6 excitatory microcircuits. This differential adenosine sensitivity may be involved in shifting the balance between subcortical and intracortical synaptic signalling.

## Method and Materials

### Slice preparation

All experimental procedures involving animals were performed in accordance with the guidelines of the Federation of European Laboratory Animal Science Association, the EU Directive 2010/63/EU, and the German animal welfare law. In this study, Wistar rats (Charles River, either sex) aged 17–21 postnatal days were obtained from Charles River and kept under a 12-h light–dark cycle, with food and water available ad libitum. Juvenile rats were housed in groups of 2-5 rats in a cage and rats older than 4 weeks were housed individually. Rats were maintained in a standard environment. Briefly, rats were deeply anaesthetised with isoflurane and decapitated. The brain was quickly removed and placed in an ice-cold artificial cerebrospinal fluid (ACSF) containing 125 mM NaCl, 2.5 mM KCl, 1.25 mM NaH2PO4, 4 mM MgCl2, 1mM CaCl2, 25mM NaHCO3, 25mM glucose, 3mM myo-inositol, 2 mM Na-pyruvate, and 0.4 mM ascorbic acid; the osmolarity of the solution was ∼310 mOsm. The extracellular Ca2+ concentration was lowered to reduce potential excitotoxic synaptic transmission during slicing. In order to maintain adequate oxygenation and a physiological pH level, the solution was constantly bubbled with carbogen gas (95% O2 and 5% CO2). 350 µm thick coronal slices of the prelimbic mPFC were cut using a vibrating micro-slicer at a low speed and high vibration frequencies. The slices were then transferred to an incubation chamber for a recovery period of ∼1 h at room temperature. Whole-cell patch-clamp recordings were made from submerged slices with continuous ACSF flowing in perfusion chamber at ∼5 ml/min. This artificial CSF containing: 125 mM NaCl, 2.5 mM KCl, 1.25 mM NaH2PO4, 1 mM MgCl2, 2 mM CaCl2, 25 mM NaHCO3, and 25 mM glucose, bubbled with carbogen gas and maintained at ∼31 ◦C. Patch pipettes were pulled from thick-wall borosilicate glass capillaries and filled with an internal solution containing 135 mM K-gluconate, 4 mM KCl, 10 mM HEPES, 10 mM phosphocreatine, 4 mM Mg-ATP, and 0.3 mM GTP (pH 7.4 with KOH, osmolarity ∼300 mOsm). Biocytin (0.5%) was added to the internal solution to obtain permanent staining of the patched neurons for morphological analysis.

### Cell identification

Slices were placed in the recording chamber under an upright microscope (equipped with 4× plan/ 0.13 numerical aperture and 40× water immersion/0.80 NA objectives; Olympus) with the pial surface facing forward. Cortical layers were distinguished by cell density and soma size, in accordance with previous studies on the PFC (Gabbott, Warner et al. 2005, van Aerde and Feldmeyer 2015, Mitric, Seewald et al. 2019, Ding, Emmenegger et al. 2021). The PFC can be divided into three sections: the upper third comprises L1–L3, the middle third L5 and the lower third L6. Putative excitatory neurons and interneurons were differentiated based on their intrinsic action potential (AP) firing pattern during recording and their morphological appearance after post-hoc histological staining. Interneurons were excluded from the analysis.

### Electrophysiology

Whole-cell patch clamp recordings were made using an EPC10 amplifier (HEKA). Signals were sampled at 10 kHz, filtered at 2.9 kHz using Patchmaster software (HEKA), and later analysed off-line using Igor Pro software (Wavemetrics). Recordings were performed using patch pipettes with a resistance of 6 to 10 MΩ. The resting membrane potential was measured immediately after establishing the whole-cell configuration. Bridge balance and capacitance neutralisation were adjusted. The whole-cell series resistance was monitored throughout the experiment and was compensated by 80%. Membrane potentials were not corrected for the junction potential. Passive and active action potential properties were assessed by injecting a series of hyper- and depolarising, current pulses under current clamp configuration for 1 s. Dual whole-cell recordings were performed in L6 of the mPFC in order to record synaptically coupled neuron pairs. Due to the low intralaminar connectivity ratio in L6, a ‘searching’ procedure was used to identify potential presynaptic neurons (Feldmeyer, Egger et al. 1999, Feldmeyer and Radnikow 2016): Putative presynaptic neurons were randomly patched in ‘loose cell-attached’ mode. When an AP elicited by a current pulse resulted in an excitatory postsynaptic potential (EPSP), this presynaptic neuron was repatched with a new pipette filled with biocytin-containing internal solution. The ‘searching’ pipette (see below) was filled with an internal solution in which K^+^ is replaced by Na^+^ (containing [in mM]: 105 Na-gluconate, 30 NaCl, 10 HEPES, 10 phosphocreatine, 4 Mg-ATP, and 0.3 GTP) in order to prevent the depolarisation of neighbouring neurons during searching for a presynaptic cell. To obtain a low staining background of pre- and postsynaptic neurons, no biocytin was added to the Na-based internal solution in ‘searching’ pipettes.

### Drug application

Adenosine (10 µM–100 µM), 8-Cyclopentyltheophylline (CPT, 1 µM), CGS21680 (30 nM), tetrodotoxin (TTX, 0.5 µM), and gabazine (10 µM) were all bath applied via the perfusion system for 3-7 minutes; drugs were purchased either from Sigma-Aldrich or Tocris.

### Histological Procedures

After single cell or paired-recordings, slices containing biocytin-filled neurons were fixed for at least 24 h at 4 °C in 100 mM phosphate buffer solution (PBS, pH 7.4) containing 4% paraformaldehyde (PFA). After several rinses in PBS, slices were treated with 1% H_2_O_2_ in PBS for approximately 20 min, in order to quench endogenous peroxidase activity. Following this, slices were rinsed several times using 100 mM PBS and subsequently incubated in 1% avidin-biotinylated horseradish peroxidase (Vector ABC staining kit, Vector Lab. Inc., Burlingame, USA) containing 0.1% Triton X-100 for 1 h at room temperature. This was followed by a chromogenic reaction that resulted in a dark precipitate by adding 0.5 mg/ml 3,3-diaminobenzidine (DAB; Sigma-Aldrich, USA) until distinct axonal and dendritic branches of the biocytin-filled neurons were clearly visible. The slices were rinsed again with 100 mM PBS, followed by slow dehydration in increasing ethanol concentrations and finally in xylene for 2–4 h. Slices were then mounted on gelatinised slides and embedded using Eukitt medium (Otto Kindler GmbH, Freiburg, Germany).

### Morphological 3D Reconstructions

Computer-assisted morphological 3D reconstructions of biocytin-filled L6 neurons were made using NEUROLUCIDA^®^ software (MicroBrightField, Williston, VT, USA) and an Olympus BV61 microscope at 1000x magnification (100x objective, 10x eyepiece). Neurons were selected for reconstruction based on the quality of biocytin labelling when background staining was minimal. The cell body, dendritic and axonal branches were reconstructed manually under constant visual inspection to detect thin and small collaterals. Layer borders, pial surface and white matter were delineated during reconstructions at lower magnification. The position of soma and layers were confirmed by superimposing the differential interference contrast images taken during the recording. The tissue shrinkage was corrected using correction factors of 1.1 in the *x*–*y* direction and 2.1 in the *z* direction (Marx, Gunter et al. 2012).

### Data analysis

#### Morphological Properties

3D reconstructed neurons were quantitatively analysed using Neuroexplorer software (MicroBrightField, Williston, VT, USA) to determine morphological properties such as the dendritic length, the number of dendritic ends, the distribution of dendritic length in cortical layers 1-6 and the white matter, and the angle of the leading dendrite. Inter-soma distances of synaptic coupled pairs was computed as the 3D-Euclidean distance between two somata. In acute brain slices, axonal collaterals, in particular those of principal cells are severely truncated during the cutting procedure (Egger, Narayanan et al. 2020). Therefore, in this study, the axonal properties of upright and inverted excitatory neurons were not included in the morphological analysis.

#### Unsupervised Hierarchical Cluster Analysis

Dendritic morphological parameters were used for unsupervised cluster analysis. Parameters were standardised using *z*-score in order to make the distributions numerically comparable. Principal component analysis (PCA) was used to analyse the interdependence between variables and to reduce the dimensionality of the dataset while preserving maximum variability. PCA reduces the redundancy of the dataset by eliminating correlated variables and produces linear combinations of the original variables to generate new axes. To determine the number of principal components to retain for cluster analysis, we used Kaiser’s rule, an objective way to determine the number of clusters by leaving all components with eigenvalues <1. Since the dataset is standardised, the variables have an eigenvalue of 1, and hence, PCs with an eigenvalue >1 describe more of the data’s variance than the original variable. Classification of excitatory neuron subtypes was then performed using unsupervised hierarchical cluster analysis employing Ward’s method (Ward 1963). Euclidean distance was used to calculate the variance. A dendrogram was constructed to visualise the distance at which clusters are combined.

#### Passive and Active Electrophysiological Properties

Custom-written macros for Igor Pro 6 (Wavemetrics) were used for the analysis of the recorded electrophysiological data. Recordings with a series resistance exceeding 45 MΩ were excluded from the data analysis. The resting membrane potential (*V*_rest_) of the neuron was measured directly after breakthrough into the whole-cell configuration with no current injection. To calculate the input resistance (*R*_in_), the slope of the linear fit to the voltage step from -60 to -70 mV of the current–voltage relationship was used. The membrane time constant was calculated as mono-exponential fit to the decay of the hyperpolarising voltage response after a current step of −50 pA. For the analysis of single spike characteristics such as threshold, amplitude, and half-width, a 10 pA step size increment for current injection was used to ensure that the action potential was elicited very close to its voltage threshold. The rheobase current was defined as the minimal current that elicited the first spike. The AP half-width was measured as the time difference between rising phase and decaying phase of the spike at half-maximum amplitude, and the AP amplitude was calculated as the difference in voltage from AP threshold to the peak during depolarization. The AP threshold was defined as the point of maximal acceleration of the membrane potential using the second derivative (d_2_*V*/d*t*^2^), *i.e.*, the time point with the fastest voltage change, and the AP latency was defined as the time required for the onset of first AP in rheobase current stimulus. The AHP amplitude was calculated as the difference in voltage from AP threshold to maximum deflection of the repolarisation. The current-frequency (If) plot was the slope of the linear fit of the current-frequency response curve between 0 and 300 pA. The interspike interval (ISI) was measured as the average time between individual spikes at a current step eliciting 10 APs. The adaptation ratio was measured as the ratio of the ninth ISI and the third ISI.

#### Synaptic Properties

Synaptic properties were evaluated as described previously (Feldmeyer, Egger et al. 1999, Feldmeyer and Radnikow 2016). All unitary EPSP (uEPSP) recordings were aligned to their corresponding presynaptic AP peaks, and an average sweep was generated as the mean uEPSP. The EPSP amplitude was calculated as the difference between the mean baseline and maximum voltage of the postsynaptic event. The paired-pulse ratio was defined as the second/third uEPSP divided by the first uEPSP amplitude elicited by presynaptic APs at a stimulation frequency of 10 Hz. Failures were defined as events with an amplitude < 1.5× the SD of the noise within the baseline window; the failure rate refers to the percentage of failures. The coefficient of variation (CV) was calculated as the SD divided by the mean uEPSP amplitude. Rise time was calculated as the mean time to rise from 20% to 80% of the peak amplitude. The latency was calculated as the time interval between the peak amplitude of presynaptic AP and the onset of the EPSP. The EPSP decay time was measured using a single exponential fit to the decay phase of both individual and averaged responses. The properties mentioned above were obtained from 20 to 80 successive sweeps.

#### Statistics

Data were either presented as box plots (n ≥ 10) or as bar histograms (n < 10). For box plots, the interquartile range (IQR) is shown as a box, the range of values that are within 1.5*IQR is shown as whiskers and the median is represent by a horizontal line in the box; for bar histograms, the mean ± SD is given. Wilcoxon Mann-Whitney U test was performed to assess the difference between individual clusters. Statistical significance was set at P < 0.05, and n indicates the number of neurons/connections analysed.

## Results

### ‘Upright’ and ‘Inverted’ excitatory neurons in layer 6 of mPFC

Excitatory neurons in layer 6 of the prefrontal cortex have generally one main leading dendrite that projects either to the pia surface or towards the white matter (WM) or both. The projection pattern of this leading dendrites exhibits a high degree of heterogeneity, which is likely to have implications for their local synaptic connectivity and network function (Zhang and Deschênes 1997, van Aerde and Feldmeyer 2015, Yang, Qi et al. 2022). Here we have classified L6 excitatory neurons mainly on the basis of the structural properties of the main dendrite with reference to their projecting angles and cortical layer distributions (**Table 1**). Principal Component Analysis (PCA) was used to analyse the interdependence between variables to avoid double weighting of the correlated variables in the cluster analysis (CA). The first three principal components (PCs) with eigenvalues larger than one were retained for CA explaining 76% of the total variance. CA based on morphological parameters grouped 79 L6 excitatory neurons into two main clusters: 48 ’upright’ neurons (61%) with main dendrites projecting to the pia, spanning superficial layers and terminating in L5 to L1; 31 inverted/ horizontally oriented neurons (39%) with a prominent main dendrite pointing to the WM or laterally oriented (**Table 1**). The two main clusters can be further subdivided into six sub-clusters (cluster 1) and two sub-clusters (cluster 2) (**Figure 1** **B, C**). The optimal number of clusters was determined using the Thorndike method, where the cut-off points have the largest linkage distance.

**Figure 1.**
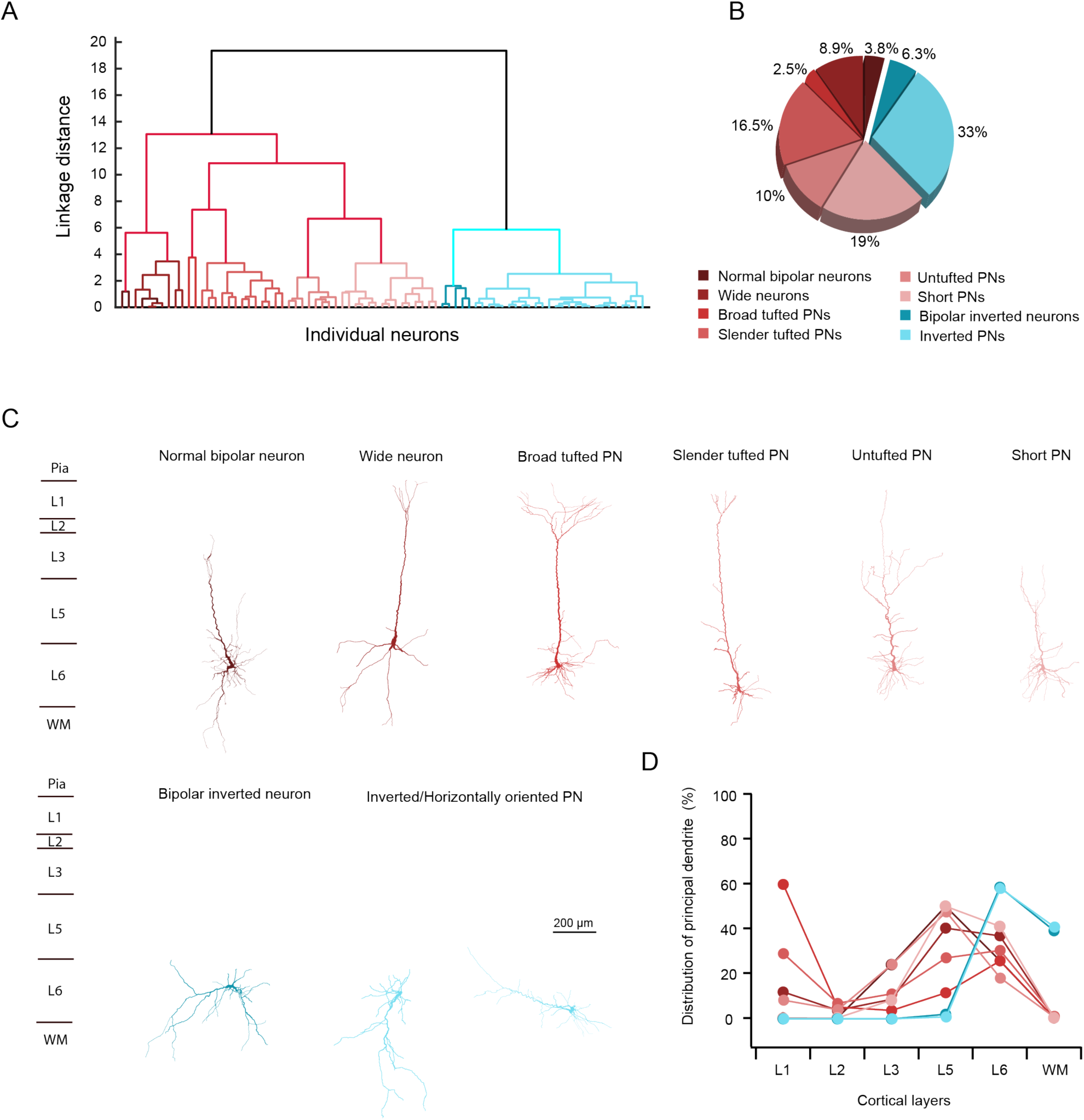
Identification of two distinct morphological types of L6 excitatory neurons in rat mPFC using cluster analysis. (**A**) Dendrogram from cluster analysis of morphological parameters reveals 2 main clusters and 8 distinct sub-clusters (n=79 neurons from 51 rats). The X-axis shows individual neurons and the Y-axis corresponds to the linkage distance measured by Euclidean distance. Cluster 1 and Cluster 2 are represented by a spectrum of colours ranging from red to blue. (**B**) Pie chart showing the percentage of each sub-cluster, with corresponding sub-cluster colours aligning with those in panel (A). (**C**) Examples of dendritic morphology from the 8 sub-clusters of L6 excitatory neurons. (**D**) The proportion of the apical dendrite extending through L1-6, and the white matter (WM) is indicated by filled dots. Each dot represents the average value derived from all neurons within the respective sub-cluster. The colour code used for the different L6 excitatory sub-clusters in panels A and B also applies to panels C and D.

**Table 1.** Morphological and electrophysiological properties of excitatory neurons in L6 of rat mPFC. Italic bold font indicates significant differences; *P < 0.05, **P < 0.01, ***P < 0.001 for Wilcoxon Mann-Whitney *U* test. Data were collected from n=51 rats.

|  | Upright neurons | Inverted neurons | Mann-Whitney Test |
| --- | --- | --- | --- |
| **Morphological properties** | **n=48 for upright and n=31 for inverted neurons** |  |  |
| **Ends of dendrite** | 18.3 ± 7.9 | 13.6 ± 4.2 | **\*\*0.0038** |
| **Length of basal dendrites (μm)** | 2412.5 ± 1084.8 | 1624.7 ± 589.8 | **\*\*\*0.0002** |
| **Length of longest basal dendrite (μm)** | 842.6 ± 589.6 | 677.7 ± 456.7 | **\*0.0436** |
| **Angle of principal dendrite** | 93.3 ± 10.2 | 260.1 ± 41.3 | **\*\*\*8.73E-011** |
| **Length of principle dendrite (μm)** | 3097.5 ± 1439.0 | 3620.7 ± 945.5 | **\*0.0436** |
| **Length-P-dendrite in L1 (μm)** | 536.3 ± 938.3 | 0.0 ± 0.0 | - |
| **Length-P-dendrite in L2 (μm)** | 103.6 ± 130.9 | 0.0 ± 0.0 | - |
| **Length-P-dendrite in L3 (μm)** | 414.6 ± 390.5 | 0.0 ± 0.0 | - |
| **Length-P-dendrite in L5 (μm)** | 1140.6 ± 538.8 | 33.2 ± 67.8 | **\*\*\*8.74E-011** |
| **Length-P-dendrite in L6 (μm)** | 905.7 ± 588.3 | 2106.2 ± 821.9 | **\*\*\*7.27E-011** |
| **Length-P-dendrite in WM (μm)** | 3.5 ± 14.9 | 1477.2 ± 828.0 | **\*\*\*3.50E-011** |
| **Electrophysiological properties** | **n=20 for upright and n=11 for inverted neurons** |  |  |
| **resting membrane potential (mV)** | -66.8 ± 5.2 | -60.5 ± 4.0 | **\*\*\*0.0006** |
| **Input resistance (MΩ)** | 192.4 ± 85.1 | 298.5 ± 117.9 | **\*\*\*4.64E-05** |
| **Membrane time constant (ms)** | 25.4 ± 5.7 | 34.3 ± 7.5 | **\*\*0.0025** |
| **Rheobase current (pA)** | 112.5 ± 39.3 | 49.1 ± 13.0 | **\*\*\*5.5E-07** |
| **AP half-width (ms)** | 1.05 ± 0.14 | 1.34 ± 0.16 | **\*\*\*7.6E-05** |
| **AP amplitude (mV)** | 88.8 ± 8.2 | 83.8 ± 6.0 | 0.1229 |
| **AP threshold (mV)** | -33.2 ± 4.5 | -29.5 ± 3.2 | **\*0.0292** |
| **AHP amplitude (mV)** | 12.5 ± 3.9 | 15.0 ± 3.2 | 0.0539 |
| **AP latency (ms)** | 246.8 ± 191.4 | 325.8 ± 223.7 | 0.2608 |
| **frequency-current slope (Hz/100 pA)** | 15.9 ± 6.8 | 20.1 ± 3.9 | **\*0.0364** |
| **ISI average (ms)** | 90.2 ± 12.8 | 78.6 ± 12.6 | **\*0.0285** |
| **Adaptation ratio (ISI<sub>3</sub>/ISI<sub>9</sub>)** | 0.86 ± 0.22 | 1.00 ± 0.30 | 0.3587 |

Of the 48 upright excitatory neurons analysed here, 3 were bipolar neurons, 7 wide pyramidal-like neurons (PNs), 2 broad tufted PNs, 13 slender tufted PNs, 8 untufted PNs and 15 short PNs. Bipolar spiny neurons have two major dendrites that are generally longer and thicker than the other dendrites and point towards the pial surface and the WM, respectively (Zhang and Deschênes 1997). Here, the longest and thickest dendrite was considered as leading or principal dendrite because of its upright projection pattern, 3 bipolar neurons were classified into the ‘upright’ neuron cluster. Broad-tufted PNs have long, sparsely tufted apical dendrites that extend as far as layer 1 and exhibit horizontally projecting basal dendrites. This dendritic morphology has been found to be typical of intratelencepalic (IT) claustrum-projecting PNs in layer 6 of the primary visual cortex (Katz 1987, Cotel, Fletcher et al. 2018). Broad-tufted PNs, slender-tufted PNs, untufted PNs and short PNs were named based on their apical dendritic distribution in different cortical layers (**Figure 1** **C, D**). Broad- and slender-tufted PNs had a tall apical dendrite that terminated close to the pial surface. A significant proportion of the apical dendritic tuft of these neurons was located in L1. The apical tuft had a large field span in the broad tufted PNs (763 ± 219 µm, n=2 neurons) but was only narrow in the slender tufted PNs (343 ± 164 µm, n=13 neurons). Untufted PNs had apical dendrites ending in either L1 or L2, but only a small proportion of the total apical dendritic length was in the terminating layer. Apical dendrites of the PNs in this sub-cluster often formed apical oblique dendrites in L3 and L5 accounting for more than 70% of the total length. Apical dendrites of short PNs terminated in L5 or L6; in general, dendrites were almost exclusively restricted to deep layers (**Figure 1** **C, D**; **Figure 1-2**).

**Figure 2.**
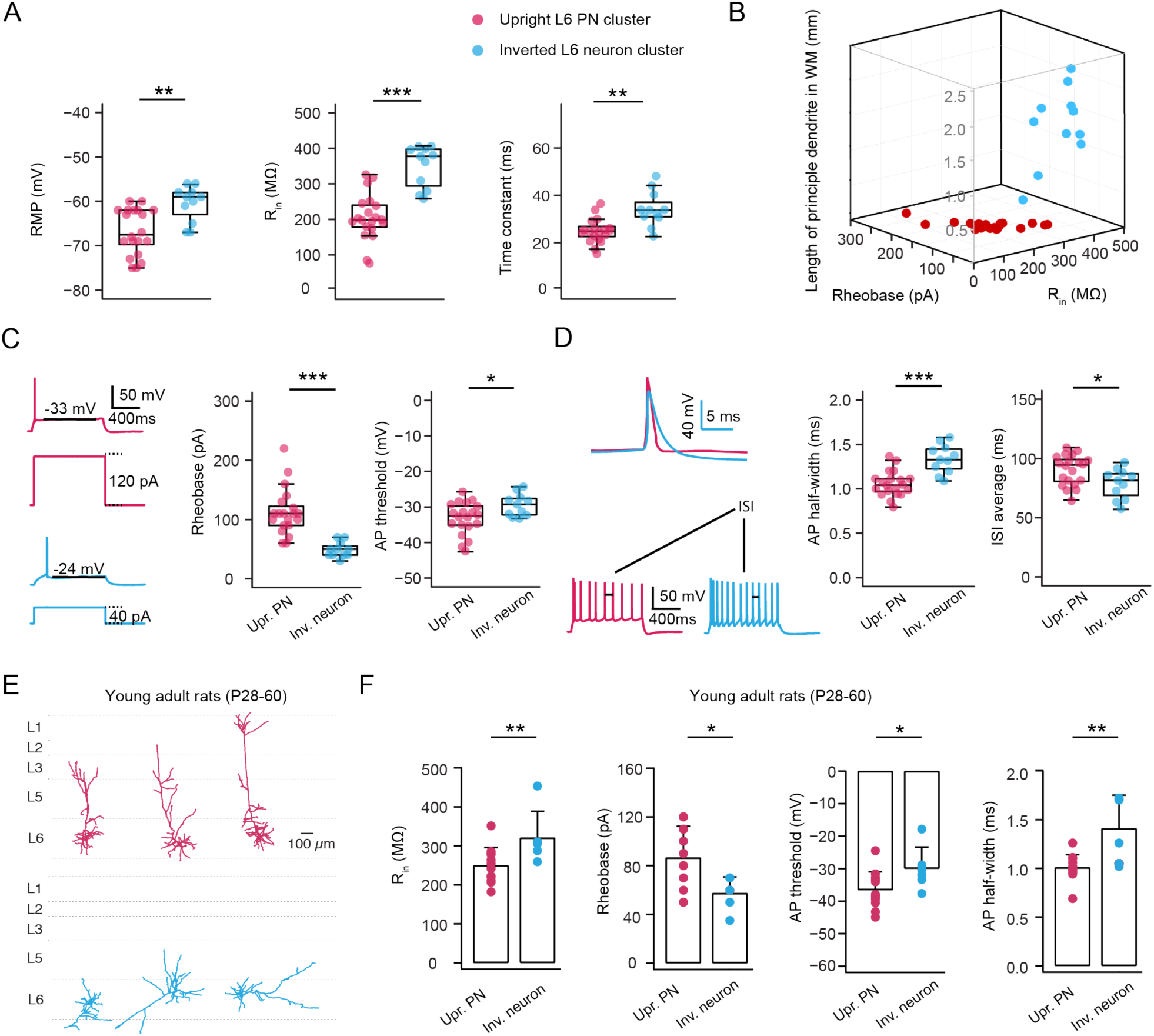
Upright and inverted L6 excitatory neurons show distinct electrophysiological properties. (**A**) Difference in resting membrane potential (RMP), input resistance (R_in_) and membrane time constant in upright L6 excitatory neurons (n=20 neurons, red) and inverted L6 excitatory neurons (n=11 neurons, cyan). **P < 0.01, ***P < 0.001 for the Wilcoxon Mann–Whitney U test. Data were obtained from 27 rats. (**B**) Difference in rheobase and AP threshold in upright L6 excitatory neurons and inverted L6 excitatory neurons. Examples are shown in the left. *P < 0.05, ***P < 0.001 for the Wilcoxon Mann–Whitney U test. (**C**) The 3D scatter plot shows a clear separation of two L6 excitatory neuron clusters using morphological and electrophysiological properties. Upright L6 excitatory neurons are shown in red and inverted L6 excitatory neurons in cyan. (**D**) Difference in AP half-width and mean inter-spike interval (ISI) in upright L6 excitatory neurons and inverted L6 excitatory neurons. Examples are shown on the left. *P < 0.05, ***P < 0.001 for the Wilcoxon Mann–Whitney U test. (**E**) Representative morphological reconstructions of L6 upright (top) and inverted (bottom) excitatory neurons recorded in mPFC of young adult rats (P28-P60). (**F**) Summary data of several electrophysiological properties of L6 upright (n = 12 neurons) and inverted (n = 6 neurons) excitatory neurons in mPFC of young adult rats (data from 8 rats).

Apart from the classical ‘upright’ excitatory neurons, there were also 31 inverted/horizontally oriented neurons including 5 bipolar inverted neurons and 26 inverted excitatory neurons. Unlike the normal bipolar neurons in cluster 1, the inverted bipolar neurons were diverse in terms of their longest secondary dendrites, but all have primary leading dendrites projecting towards the white matter. In the sub-cluster of inverted excitatory neurons, neurons have only one major dendrite projecting either towards and often into the WM or horizontally to it. All neurons in cluster 2 have main dendrites restricted to L6 and WM (**Figure 1** **C, D**). Notably, the leading dendrites of L6 inverted neurons are longer than those of upright neurons (3621 ± 946 vs. 3098 ± 1439 µm, *P < 0.05), but the total length of basal dendrites is shorter (1625 ± 590 vs. 2413 ± 1085 µm, ***P < 0.001).

### Upright and inverted excitatory neurons show distinct electrophysiological properties

To determine whether neurons belonging to either of the two morphological clusters identified here have different electrophysiological properties, their passive membrane properties, single AP properties and firing behaviour were analysed. Neurons with a resting membrane potential less negative than -55 mV or a series resistance higher than 40 MΩ were excluded from the final data set. Therefore, electrophysiological parameters from 20 upright neurons and 11 inverted neurons were used for statistical comparison.

L6 upright and inverted excitatory neurons show significant differences in their passive and active electrophysiological properties. Inverted excitatory neurons had a more depolarised resting membrane potential (-60.55 ± 4.03 vs. -66.8 ± 5.18 mV, ** P < 0.01) when compared to upright excitatory neurons. Although the input resistance (R_in_) of neurons in both clusters was higher than that of neurons in other layers (van Aerde and Feldmeyer 2015), it was exceptionally high in inverted excitatory neurons, whose R_in_ was almost twice that of upright excitatory neurons, (396.08 ± 95.97 vs. 235.93 ± 77.26 MΩ, ***P< 0.001), resulting in a slower membrane time constant (34.27±7.46 vs. 25.35±5.66 ms, P< 0.001) for inverted neurons (**Figure 2A**). This indicates that the two L6 neuron clusters differed significantly in their excitability. Consistent with this, upright neurons were significantly less excitable than inverted neurons as indicated by rheobase values of 112.5 ± 39.32 and 49.09 ± 13 pA, respectively (***P< 0.001). Furthermore, upright excitatory neurons showed a more hyperpolarised AP threshold compared to inverted neurons (-29.46 ± 3.16 vs. -33.21 ± 4.54 mV, *P < 0.05; **Figure 2B**). The 3D scatter plot in **Figure 2C** shows that excitatory neurons in layer 6 of the mPFC can be reliably distinguished by both morphological and physiological features.

In addition, the AP half-width was much shorter in L6 upright excitatory neurons than in inverted excitatory neurons (1.34 ± 0.16 vs. 1.06 ± 0.14 ms, ***P < 0.001), as expected from the higher capacitive load (i.e. high R_m_ and slow τ_m_) in the latter. Most L6 excitatory neurons in the mPFC had a regular firing pattern; however, some of these regular spiking neurons displayed a very short first inter-spike interval (ISI) at the beginning of the spike train. As this affects the second ISI, the adaptation ratios were measured as the ratio of third and ninth ISI (ISI_3_/ISI_9_) for all excitatory neurons. There was no major difference in AP firing adaptation of both upright and inverted excitatory neurons. Using current injections of the same amplitude, inverted excitatory neurons fired more APs, resulting in a significantly smaller mean ISI value than upright excitatory neurons (78.63 ± 12.6 vs. 90.19 ± 12.78 ms, *P < 0.05). Further electrophysiological properties and the statistical comparison of the two excitatory L6 neuron types are shown in **Table 1**.

It has been reported that adenosine-mediated modulation of synaptic transmission and plasticity displays an ontogenic variation (Descombes, Avoli et al. 1998, Costenla, Diogenes et al. 2011). To investigate the variation of neuronal characteristics in the two distinct groups during postnatal development, we conducted whole-cell recordings from L6 excitatory neurons in rats aged between 4 weeks and 2 months. Among 18 recorded neurons, 12 are upright and 6 are inverted excitatory neurons, reflecting a ratio of 67% to 33%, which is similar to our findings of two morphological groups in juvenile rats (61% to 39%). Electrophysiological properties were compared between upright and inverted neurons. Consistent with our findings in juvenile rats, inverted neurons exhibited higher input resistance, a smaller rheobase current, a more depolarized action potential (AP) threshold, and a broader AP half-width (**Figure 2E,F**). These observations suggest that the differences in electrophysiological properties between upright and inverted excitatory neurons persist from pre-adolescence into young adulthood.

### Upright and inverted neurons form synaptic connections with distinct functional and structural properties

Due to the low excitatory synaptic connectivity ratio in layer 6, a so-called ‘loose seal’ search protocol (see ‘Methods and Materials’) was used to test for potential synaptic connections. A total of 59 neurons were recorded as postsynaptic neurons. Out of 1450 potential presynaptic neurons, 45 neurons were found to be connected to the recorded postsynaptic neurons, i.e. the connectivity ratio was 3.1% in layer 6 of the mPFC. After re-patching, 20 excitatory monosynaptic connections were recorded in dual whole-cell mode. In 15 out of 20 pairs the pre- and postsynaptic neurons were morphologically reconstructed and classified according to their dendritic morphology as described in **Figure 1**. We identified 7 synaptically coupled pairs with a presynaptic upright neuron and 8 pairs with a presynaptic inverted excitatory neuron. All presynaptic upright neurons were synaptically coupled to another postsynaptic upright excitatory neuron whereas inverted neurons formed connections with either an upright (n=3) or an inverted (n=5) neuron.

The mean inter-soma distance was found to be significantly shorter in synaptic connections between two upright excitatory neurons compared to those with a presynaptic inverted neuron (61.4 ± 33.0 vs. 107.7 ± 32.0, *P < 0.05) (**Figure 3A**). The E→E connections with a presynaptic upright excitatory neurons displayed unitary EPSPs (uEPSPs) with an average amplitude of 0.22 ± 0.23 mV, which is significantly smaller than that evoked by presynaptic inverted excitatory neurons (1.03 ± 0.61 mV, *P < 0.05). Most pairs (6 out of 7) with a presynaptic upright excitatory neuron showed strong short-term facilitation resulting in an average paired-pulse ratio (PPR) of 3.6 ± 3.9. In contrast, connections established by inverted excitatory neurons all showed short-term depression, characterised by a mean PPR of 0.68 ± 0.14 (**Figure 3B**).

**Figure 3.**
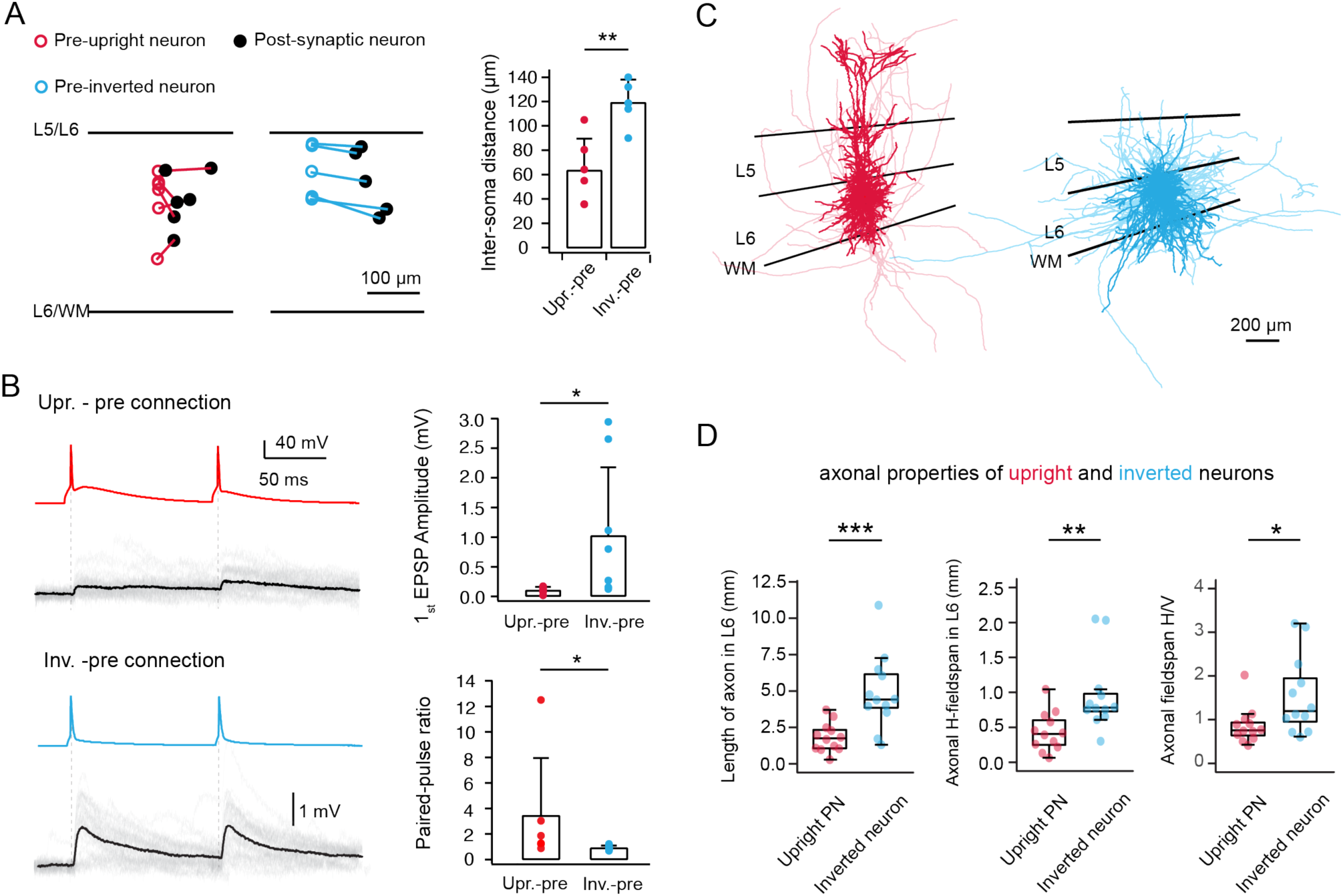
Functional properties of L6 E*⟶*E connections depend on the different presynaptic neuron subtypes. **(A)** Left, the position in layer 6 and inter-soma distance of pre- and postsynaptic neurons in each pair are shown. Presynaptic neurons are colour-coded in red and cyan for upright and inverted neurons, respectively; postsynaptic neurons are shown in black. The plot was constructed by aligning the L5/L6 borders. Right, Histogram comparing the inter-soma distance for connections with a presynaptic upright and presynaptic inverted L6 neuron, respectively. **P < 0.01 for the Wilcoxon Mann–Whitney U test. **(B)** Left, unitary synaptic connections in mPFC layer 6 with a presynaptic upright (top) or inverted (bottom) excitatory neuron. EPSPs are elicited by two presynaptic APs at an interstimulus interval of 100 ms. The mean uEPSP waveform is shown in black and the individual sweeps are shown in light grey. Right, histograms showing differences in first uEPSP amplitude and paired pulse ratio between upright- (n=7) and inverted-pre (n=8) connections. **P < 0.05 for the Wilcoxon Mann–Whitney U test (data from 15 rats). **(C)** Superimposed morphological reconstructions of upright (red, n=15) and inverted (cyan, n=15) neurons. The somatodendritic domain is shown in a darker, the axons in a lighter shades. Layer borders of L3/L5, L5/L6 and L6/WM are indicated by black lines from top to bottom (data from 17 rats). **(D)** Summary data of several axonal properties of L6 upright (n = 12) and inverted (n = 12) excitatory neurons in rat mPFC (data from 14 rats).

It has been reported that inverted pyramidal neurons have long and dense axonal projections in deep neocortical layers, thereby contributing to a network of long-range intracortical connections between different cortical areas (Zhang and Deschênes 1997, Egger, Narayanan et al. 2020). To investigate whether the different synaptic properties depend on the distinct axonal morphology of these two L6 neuron types, we reconstructed the axons of L5 neurons in each morphology cluster. We found that the length of axon located in layer 6 was found to be significantly smaller for upright excitatory neurons than inverted excitatory neurons (2239 ± 1319 vs. 4553 ± 2463 µm, **P < 0.01). Furthermore, the absolute horizontal axonal field span in layer 6 of upright L6 excitatory neurons was significantly smaller than that of inverted L6 excitatory neurons (491 ± 273 vs. 906 ± 492 µm, **P < 0.01). The ratio between the horizontal and vertical axonal field span was also significantly different between upright and inverted excitatory neurons (0.92 ± 0.42 and 1.56 ± 0.81, respectively; *P < 0.05), indicating that inverted excitatory neurons have a prominent axonal projections along the horizontal axis (**Figure 3** **C, D**). It should be noted that for inverted neurons, this ratio is severely underestimated due to the high degree of truncation of long-range axonal collaterals during acute brain slice preparation. Taken together, our results demonstrate a strong correlation between presynaptic neuronal morphology and postsynaptic uEPSP properties.

### The neuromodulator adenosine reduces presynaptic neurotransmitter release in L6 excitatory neurons via activating A_1_ARs

To elucidate the adenosinergic modulation of excitatory synaptic transmission, paired recordings were performed from synaptically coupled mPFC L6 neurons (**Figure 4A**). We found that adenosine strongly suppressed the synaptic efficacy of all the L6 excitatory connections without exception (**Figure 4B-D**). For L6 excitatory connections, the postsynaptic mean uEPSP amplitude decreased from 0.75 ± 0.67 mV to 0.14 ± 0.15 mV (n=15 connections, ***P< 0.001) while the PPR increased from 1.12 ± 0.80 to 2.42 ± 2.45 (***P< 0.001) after bath application of 30 µM adenosine. In addition, the CV increased from 0.82 ± 0.47 to 1.29 ± 0.33 (**P< 0.01) and the failure rate increased from 31.9 ± 31.4 % to 72.3 ± 16.2% (***P< 0.001) at these connections (**Figure 4D**). No significant changes in EPSP latency, 20-80% rise time and decay time were found, suggesting that adenosine suppresses synaptic transmission of excitatory connections predominantly via a presynaptic mechanism (**Table 2**). To investigate whether adenosine inhibits presynaptic release in a cell type-dependent manner, presynaptic neurons were classified either as upright or inverted neurons based on the above criteria. We found that bath application of 30 µM adenosine reduced the amplitude of uEPSP at connections with a presynaptic upright neuron to 27 ± 16 % of control, whereas at connections with a presynaptic upright excitatory neuron the uEPSP amplitude was reduced to 25 ± 21 % of control. Thus, following application of 30 µM adenosine, no significant change in the percentage of EPSP amplitude reduction was found between these two types of excitatory connections (**Figure 4D**).

**Figure 4.**
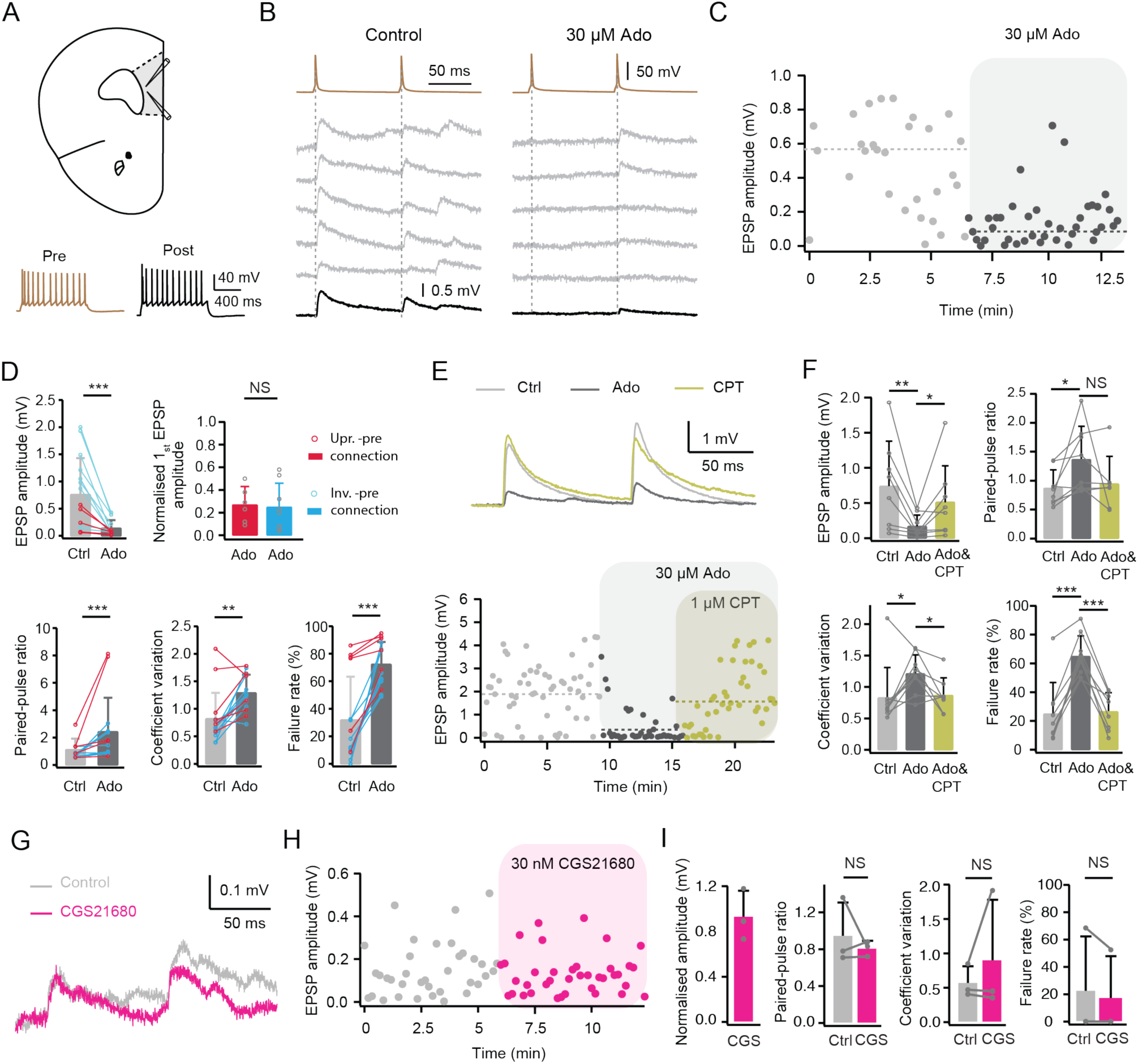
Adenosine suppresses presynaptic release probability of excitatory connections in layer 6 of the mPFC via A_1_ARs. **(A)** Top, schematic representation of a mPFC slice with the position of the paired-recording electrodes. Bottom, pre- and postsynaptic firing patterns are shown in in brown and black, respectively. **(B)** Current clamp recordings of unitary synaptic connections in layer 6 of rat mPFC. Five consecutive uEPSPs (middle, light grey) and the average EPSP (bottom, black) are elicited by two presynaptic APs at 10 Hz (top, brown). **(C)** Time course of EPSP amplitude change following bath application of 30 µM adenosine to a L6 excitatory connection. Dashed lines indicate the mean first EPSP amplitude under control and adenosine conditions. **(D)** Summary data of several EPSP properties for L6 excitatory connections are shown. Error bars represent SD. NS (not significant), **P < 0.01, ***P < 0.001 for Wilcoxon signed-rank test. n=15 pairs from 15 rats. **(E)** Top, superimposition of EPSPs recorded in control, the presence of Ado (30 µM), and of Ado and CPT (1 µM) from a representative excitatory connection in layer 6 of rat mPFC. Bottom, time course of EPSP amplitude changes following bath application of 30 µM adenosine (ADO) and of adenosine and CPT (ADO&CPT) in the same excitatory pair. **(F)** Histograms (n=9 pairs from 9 rats) showing the effect of adenosine (ADO) and CPT blockade of adenosine-induced changes (ADO&CPT) on several EPSP properties including EPSP amplitude, PPR, CV, and failure rate. Error bars represent SD. *P < 0.05, **P < 0.01, ***P < 0.001 for Wilcoxon signed-rank test. **(G)** Superimposition of EPSPs recorded in control and the presence of CGS21680 (30 nM) from a representative excitatory connection in layer 6 of rat mPFC. **(H)** Time course of EPSP amplitude change during application of CGS21680. **(I)** Summary data of CGS21680-induced changes in EPSP properties (n=3 pairs from 3 rats). NS, not significant for paired t-test.

**Table 2.**
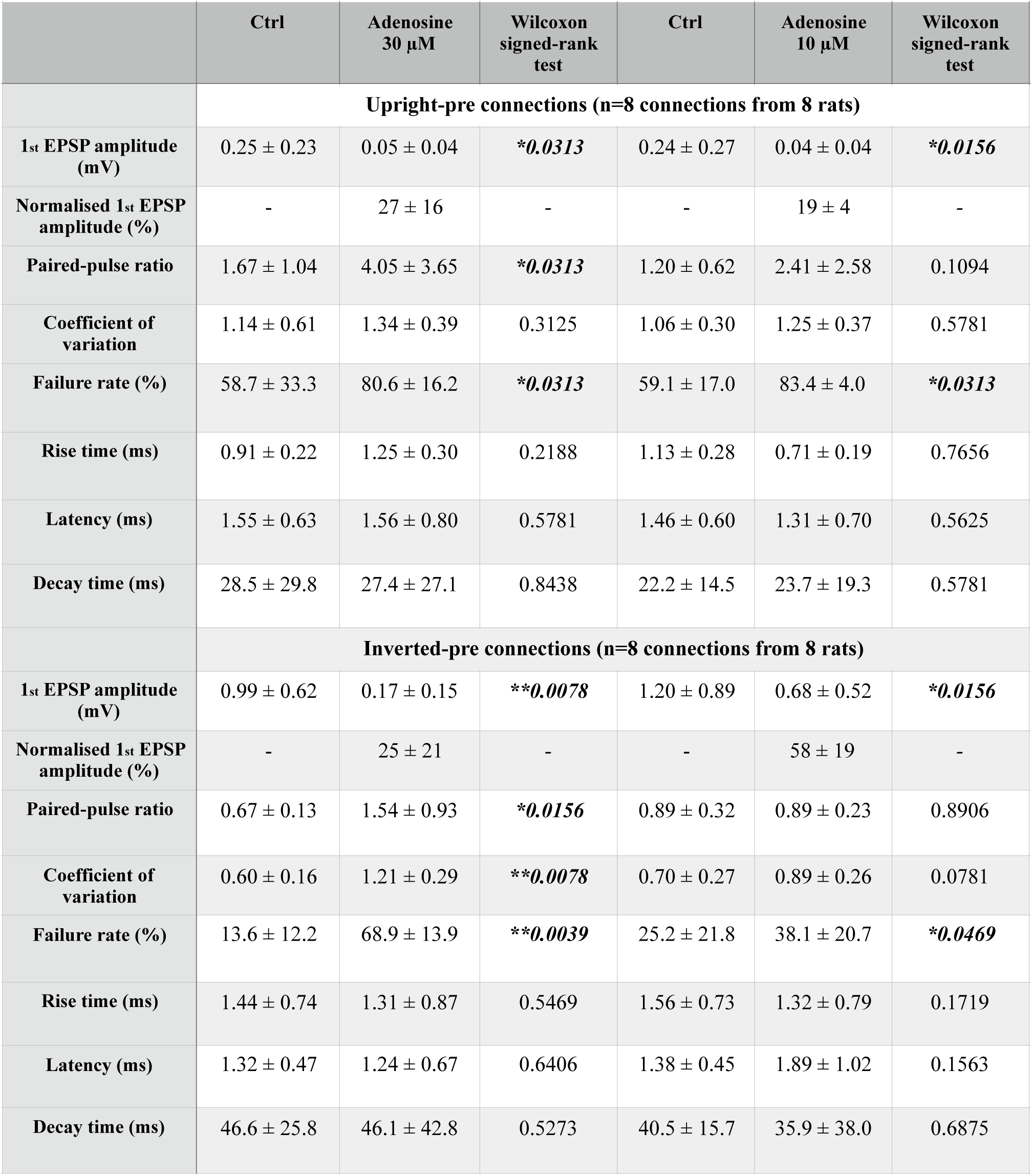
uEPSP properties of L6 upright- and inverted-pre connections under control and adenosine conditions. Bold italic font indicates significant differences to control; *P < 0.05, **P < 0.01, ***P < 0.001 for Wilcoxon signed-rank test.

| | Ctrl | Adenosine<br>30 μM | Wilcoxon<br>signed-rank<br>test | Ctrl | Adenosine<br>10 μM | Wilcoxon<br>signed-rank<br>test |
| --- | --- | --- | --- | --- | --- | --- |
|  | **Upright-pre connections (n=8 connections from 8 rats)** |  |  |  |  |  |
| **1<sup>st</sup> EPSP amplitude (mV)** | 0.25 ± 0.23 | 0.05 ± 0.04 | ***\*0.0313*** | 0.24 ± 0.27 | 0.04 ± 0.04 | ***\*0.0156*** |
| **Normalised 1<sup>st</sup> EPSP amplitude (%)** | - | 27 ± 16 | - | - | 19 ± 4 | - |
| **Paired-pulse ratio** | 1.67 ± 1.04 | 4.05 ± 3.65 | ***\*0.0313*** | 1.20 ± 0.62 | 2.41 ± 2.58 | 0.1094 |
| **Coefficient of variation** | 1.14 ± 0.61 | 1.34 ± 0.39 | 0.3125 | 1.06 ± 0.30 | 1.25 ± 0.37 | 0.5781 |
| **Failure rate (%)** | 58.7 ± 33.3 | 80.6 ± 16.2 | ***\*0.0313*** | 59.1 ± 17.0 | 83.4 ± 4.0 | ***\*0.0313*** |
| **Rise time (ms)** | 0.91 ± 0.22 | 1.25 ± 0.30 | 0.2188 | 1.13 ± 0.28 | 0.71 ± 0.19 | 0.7656 |
| **Latency (ms)** | 1.55 ± 0.63 | 1.56 ± 0.80 | 0.5781 | 1.46 ± 0.60 | 1.31 ± 0.70 | 0.5625 |
| **Decay time (ms)** | 28.5 ± 29.8 | 27.4 ± 27.1 | 0.8438 | 22.2 ± 14.5 | 23.7 ± 19.3 | 0.5781 |
|  | **Inverted-pre connections (n=8 connections from 8 rats)** |  |  |  |  |  |
| **1<sup>st</sup> EPSP amplitude (mV)** | 0.99 ± 0.62 | 0.17 ± 0.15 | ***\*\*0.0078*** | 1.20 ± 0.89 | 0.68 ± 0.52 | ***\*0.0156*** |
| **Normalised 1<sup>st</sup> EPSP amplitude (%)** | - | 25 ± 21 | - | - | 58 ± 19 | - |
| **Paired-pulse ratio** | 0.67 ± 0.13 | 1.54 ± 0.93 | ***\*0.0156*** | 0.89 ± 0.32 | 0.89 ± 0.23 | 0.8906 |
| **Coefficient of variation** | 0.60 ± 0.16 | 1.21 ± 0.29 | ***\*\*0.0078*** | 0.70 ± 0.27 | 0.89 ± 0.26 | 0.0781 |
| **Failure rate (%)** | 13.6 ± 12.2 | 68.9 ± 13.9 | ***\*\*0.0039*** | 25.2 ± 21.8 | 38.1 ± 20.7 | ***\*0.0469*** |
| **Rise time (ms)** | 1.44 ± 0.74 | 1.31 ± 0.87 | 0.5469 | 1.56 ± 0.73 | 1.32 ± 0.79 | 0.1719 |
| **Latency (ms)** | 1.32 ± 0.47 | 1.24 ± 0.67 | 0.6406 | 1.38 ± 0.45 | 1.89 ± 1.02 | 0.1563 |
| **Decay time (ms)** | 46.6 ± 25.8 | 46.1 ± 42.8 | 0.5273 | 40.5 ± 15.7 | 35.9 ± 38.0 | 0.6875 |

The inhibitory effects of adenosine on synaptic transmission via A_1_ARs have been extensively investigated in the hippocampus (Dunwiddie and Hoffer 1980, Dunwiddie and Fredholm 1989, Moore, Nicoll et al. 2003, Hargus, Bertram et al. 2009). Previous studies have suggested that presynaptic A_1_ARs are responsible for modulating excitatory synaptic transmission in upper layers of the piriform, entorhinal and somatosensory barrel cortices (Yang, Chiu et al. 2007, Wang, Kurada et al. 2013, Qi, van Aerde et al. 2017). To investigate whether the effect of adenosine on synaptic transmission in layer 6 is induced by A_1_AR activation, the A_1_AR antagonist CPT (1 µM) was co-applied with adenosine (30 µM) after adenosine had been applied alone. The effect of adenosine on excitatory connections was blocked and recovered from 0.21± 0.18 mV to 0.52 ± 0.50 mV of the control level (n=9 connections, *P< 0.05). Co-application of adenosine and CPT also blocked the adenosine-induced changes in other EPSP properties including the CV and failure rate (**Figure 4E,F**). This suggest that adenosine decreases the neurotransmitter release probability via activation of presynaptic A_1_ARs.

In L2/3 of rat visual cortex, A_2A_ARs have no direct effect on synaptic transmission, but can modulate its A_1_AR-mediated suppression (Bannon, Zhang et al. 2014, Zhang, Bannon et al. 2015). In L5 of rat mPFC, A_2A_R modulate long-term plasticity at excitatory synapses onto fast-spiking interneurons rather than synaptic transmission between glutamatergic neurons (Kerkhofs, Canas et al. 2018). To elucidate whether A_2A_ARs also participate in the adenosine-induced effects on synaptic transmission in L6 of mPFC, 30 nM of the specific A_2A_AR agonist CGS21680 was bath-applied during recordings of excitatory connections. Synaptic properties correlated with neurotransmitter release probability were not altered by CGS21680 at L6 excitatory connections (**Figure 4G-I**). Single cell recordings showed no change in the postsynaptic resting membrane potential following bath application of CGS21680 suggesting that the A_2A_AR expression in mPFC L6 excitatory neurons is low at least at the soma.

To further test the presynaptic effect of adenosine, miniature spontaneous postsynaptic activity of L6 excitatory neurons was recorded in voltage-clamp mode. In the presence of 0.5 µM TTX and 10 µM gabazine in the bath, miniature excitatory postsynaptic currents (mEPSCs) were recorded under control conditions and in the presence of adenosine (**Figure 5A**). Bath application of 30 µM adenosine significantly reduced the frequency of mEPSCs from 1.0 ± 0.5 to 0.6 ± 0.4 Hz (n=12 neurons, ***P< 0.001), suggesting that an AP-independent mechanism is also involved in adenosine-induced inhibition of glutamatergic transmission. No significant change in the mEPSC amplitude was observed (10.9 ± 2.1 pA for control vs. 10.8 ± 2.8 pA for adenosine) (**Figure 5B, C**). Co-application of adenosine and CPT completely blocked the adenosine-induced decrease of the mEPSCs frequency (0.99 ± 0.52 for control vs. 0.90 ± 0.43 for adenosine & CPT, n=15 neurons, *P* = 0.43), indicating that decrease in mEPSC frequency in mPFC L6 excitatory neurons is mediated by A_1_ARs activation (**Figure 5D, E**). To reveal the effect of endogenous adenosine on L6 neurons, CPT was applied in the absence of adenosine. CPT significantly increased the miniature excitatory postsynaptic current (mEPSC) frequency in L6 neurons from 0.99 ± 0.52 to 1.20 ± 0.65 Hz (**Figure 5** **F&G**). Comparison of the relative proportion of this increase between the two neuronal clusters did not yield any statistically significant differences, indicating that endogenous adenosine exerts an inhibitory effect on presynaptic inputs on L6 excitatory neurons.

**Figure 5.**
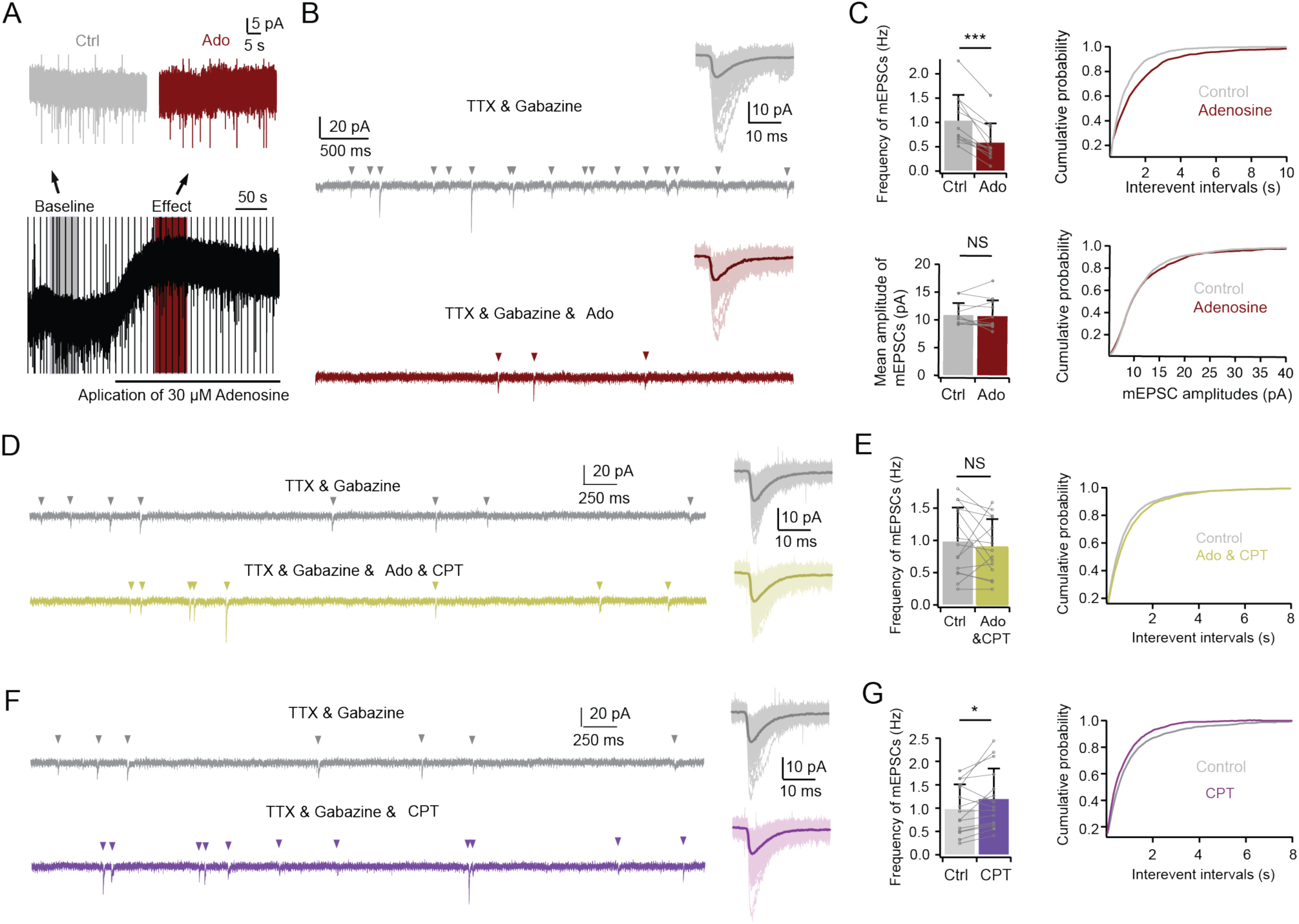
Modulation of miniature spontaneous synaptic activity in L6 excitatory neurons by adenosine through A_1_ receptor activation. **(A)** Representative voltage-clamp recording illustrating spontaneous activity during application of TTX (0.5 µM), gabazine (10 µM), and adenosine (30 µM). **(B)** Comparison of 5 s recording epochs under control (grey) and norepinephrine (NE) conditions (dark red). Insets show overlay of excitatory postsynaptic currents (EPSCs) extracted from a 50-second continuous recording depicted in panel (A). The average and individual EPSCs are superimposed and given in a darker and lighter shade, respectively. **(C)** Adenosine application (30 µM) led to an increase in the inter-event interval (IEI, right, 1114 samples for control and 626 samples for adenosine from 12 cells) and a decrease in the frequency of miniature excitatory postsynaptic currents (mEPSCs), without significant alteration in mEPSC amplitudes (n=7 for upright and n=5 for inverted neurons; data from 5 rats). NS (not significant), ***P < 0.001 for Wilcoxon signed-rank test. Error bars represent SD. **(D)** Comparative 5 s recording epochs under control (grey) and co-application of adenosine (30 µM) and 8-cyclopentyltheophylline (CPT, 1 µM) conditions (light green). Insets display overlay of EPSCs extracted from 50-second continuous recordings under control and pharmacological conditions. The average and individual EPSCs are superimposed and given in a darker and lighter shade, respectively. **(E)** Co-application of adenosine (30 µM) and CPT (1 µM) restored adenosine-induced reduction in mEPSC frequency to control levels (n=15 neurons; data from 7 rats). NS (not significant) for Wilcoxon signed-rank test. Error bars represent SD. **(F)** Comparative 5-second recordings under control (grey) and application of CPT (1 µM) conditions (purple). Insets display overlay of EPSCs extracted from 50-second continuous recordings under control and pharmacological conditions. The average and individual EPSCs are superimposed and given in a darker and lighter shade, respectively. **(G)** Application of CPT alone slightly increased mEPSC frequency in both upright (n=11) and inverted excitatory neurons (n=6; data from 7 rats). NS (not significant), *P < 0.05 for Wilcoxon signed-rank test. Error bars represent SD.

### Connections with a presynaptic upright excitatory neuron are more sensitive to low concentration of adenosine

In the PFC, the concentration of endogenous adenosine release ranges widely from nanomolar to micromolar concentration (Fredholm 2007, Nguyen, Lee et al. 2014). In layer 4 of the barrel cortex, adenosine suppresses excitatory synaptic transmission with a half-maximal effective concentration of 9.6 µM (Qi, van Aerde et al. 2017). Here, we used a lower concentration of adenosine (10 µM) to test whether presynaptic upright and inverted neurons show differential sensitivity to adenosine. Although the connections established by an inverted excitatory neuron and an upright excitatory neuron showed no significant difference in their response to 30 µM adenosine, we found that connections established by L6 upright excitatory neurons are more sensitive to a lower concentration of adenosine. Bath application of 10 µM adenosine decreased the uEPSP amplitude at connections with a presynaptic upright excitatory neuron to 19 ± 4 % of control (**Figure 6A, C-D**). In contrast, the uEPSP amplitude at connections with a presynaptic inverted excitatory neuron was reduced to only 58 ± 19 % of control after applying the same adenosine concentration (**Figure 6B, E-F**). This was significantly higher compared to the connections established by an upright excitatory neurons (n=8 for each connection type, ***P < 0.001, **Figure 6G**, **Table 2**). More details of the functional properties of two connection types under control and 10 µM adenosine conditions are shown in **Table 2**.

**Figure 6.**
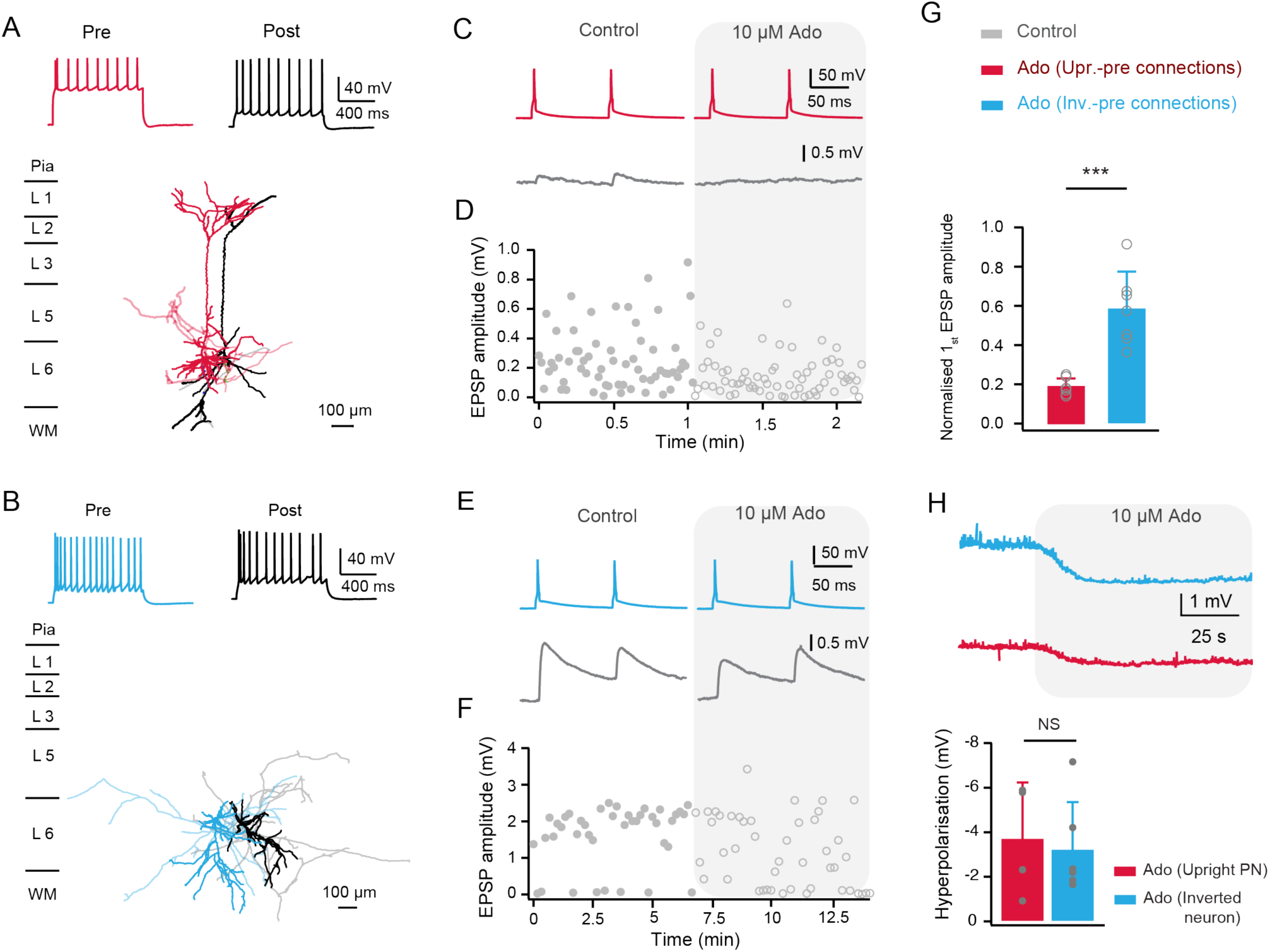
Excitatory connections with a presynaptic upright or inverted neuron showed different sensitivities to adenosine. **(A, B)** Morphological reconstructions of a synaptically coupled L6 upright-upright (A) and inverted-inverted pair. The presynaptic somatodendritic domain is shown in a darker red and blue for upright and inverted neurons, respectively; their axons are shown light red and blue. Postsynaptic soma and dendrites are in black, and postsynaptic axons are in grey. The corresponding firing patterns of pre- and postsynaptic neurons are shown on the top. **(C, E)** Average EPSPs recorded in control and the presence of adenosine (10 µM) from a representative connection with a presynaptic upright (C) and an inverted (E) excitatory neuron. Presynaptic APs are shown at the top. **(D, F)** Time course of EPSP amplitude changes during application of 10 µM adenosine for a connection with a presynaptic upright (D) and inverted (F) neuron. **(G)** Summary data (connections with presynaptic upright neurons, n=7; connections with presynaptic inverted neurons, n=7) of adenosine-induced changes in EPSP amplitude. Data were collected from 14 rats. ***P < 0.001 for Wilcoxon Mann–Whitney U test, error bars represent SD. **(H)** Top, representative current-clamp recordings of a postsynaptic upright and inverted neuron following bath application of 10 µM adenosine. Bottom, Adenosine-induced (10 µM) hyperpolarisation of the resting membrane potential with two neuron types (inverted excitatory neurons, n=6, upright excitatory neurons, n=5; data from 11 rats). NS, not significant for Wilcoxon Mann–Whitney U test, error bars represent SD.

To test whether the difference in adenosine sensitivity was mediated by postsynaptic adenosine receptors, the postsynaptic membrane potential was measured simultaneously following bath application of 10 µM adenosine. No significant difference in the amplitude of adenosine-induced hyperpolarisation was found between the two types of L6 excitatory neuron (**Figure 6H**). These findings suggest that the presynaptic release sites of upright excitatory neurons have a greater sensitivity to low concentration of adenosine compared to those of inverted excitatory neurons. Nevertheless, the change in membrane potential is predominantly mediated by receptors located at the soma; therefore it is possible that the two neuron types also differ in their adenosine sensitivity at the postsynaptic dendritic sites.

## Discussion

Layer 6 of the medial prefrontal cortex (mPFC) remains a largely unexplored brain region with a highly heterogeneous cellular composition. Here we classified excitatory neurons in mPFC layer 6 into two main morphological groups based on their dendritic arborisation pattern: upright pyramidal cell-like and inverted neurons. We found that 1) upright and inverted excitatory neurons have different electrophysiological properties with the latter having a higher intrinsic excitability. 2) Upright excitatory neurons establish weak, facilitating synapses with other L6 neurons whereas synapses formed by inverted cells show a higher EPSP amplitude and a pronounced short-term depression. This may be due to the distinct axonal projecting pattern of these two L6 neuron types within their home layer. In addition, 3) adenosine induced a reduction of the neurotransmitter release probability in L6 excitatory neurons via activation of presynaptic A_1_ARs. Although 30 µM adenosine strongly suppressed connections with both presynaptic upright or inverted excitatory neurons, 4) synapses formed by L6 upright neurons are more sensitive to low concentrations of adenosine (10 µM). Our results reveal that two functionally and morphologically distinct subpopulations of L6 principal neurons are differentially affected by adenosine, suggesting that they may play distinct roles in the maintenance of sleep homoeostasis by adenosine.

### Upright and inverted projection neurons in layer 6 of mPFC

L6 excitatory neurons show a high degree of morphological diversity in all neocortical areas including the mPFC (Yang, Seamans et al. 1996, Zhang and Deschênes 1997, Kumar and Ohana 2008, Thomson 2010, Pichon, Nikonenko et al. 2012, Marx and Feldmeyer 2013, van Aerde and Feldmeyer 2015, Yang, Qi et al. 2022). In the neocortex the incidence of inverted excitatory neurons is as low as 1% of rat (Parnavelas, Lieberman et al. 1977) and this fraction varies through species and brain area (up to 8.5%) (Globus and Scheibel 1967, Qi, Jain et al. 1999, Mendizabal-Zubiaga, Reblet et al. 2007). They are most numerous in deep layers in all studied species (Mendizabal-Zubiaga, Reblet et al. 2007). In rat barrel cortex the percentage of inverted neurons among all excitatory neurons in layer 6A was only 2.6% (Yang, Qi et al. 2022) but was found to be up to 10% in layer 6B (Marx and Feldmeyer 2013). Due to the small proportion of inverted spiny cells in the total principal cell population, their electrophysiological and functional properties have not been systematically investigated. It is worth noting that inverted excitatory neurons were found in abundance in reeler mutant mice which are characterised by an aberrant cortical layer formation due to reelin deficiency during the corticogenesis (Landrieu and Goffinet 1981, Forster 2014). In this study, we conducted an unsupervised cluster analysis (CA) based on neuronal dendritic morphology. This approach allowed for an objective and quantitative classification of mPFC L6 excitatory neurons, grouping them into distinct categories based on their leading dendritic orientation: upright or inverted/horizontally oriented neurons. Our findings revealed that non-pyramidal neurons constituted a substantial portion, accounting for 39% of the total principal neuron populations. Even when excluding bipolar inverted (n=5) and horizontally oriented (n=1) neurons, 25 typical inverted excitatory neurons remained, comprising 32% of our sample of L6 excitatory neuron. This remarkable prevalence of inverted excitatory neurons highlights their potentially crucial role within the L6 microcircuitry of the mPFC.

In addition to the distinct dendritic morphology, further differences were observed when comparing the electrophysiological properties of the two morphologically identified types of mPFC L6 excitatory neurons. Although all the excitatory neurons in layer 6 are unique with their high cellular input resistance resulting in a high excitability (van Aerde and Feldmeyer 2015), inverted excitatory neurons have a more depolarised resting membrane potential and AP threshold, larger membrane input resistance and time constant, longer AP half-width, smaller rheobase when compared to upright excitatory neurons. This is in consistent with a previous study in primary somatosensory cortex, which reported that inverted excitatory neurons have different passive and active physiological properties compared to upright excitatory neurons (Steger, Ramos et al. 2013). Differences between inverted and upright excitatory neurons in resting membrane potential and input resistance suggest a potential difference in the distribution of ion channels that determine intrinsic electrophysiological properties, such as the potassium leak channel (Lesage 2003). Differential AP threshold, halfwidth, and firing frequency are probably due to differences in ion channels contributing to action potential initiation and repolarisation such as voltage-gated sodium channel and potassium channel types (Miller, Okaty et al. 2008). Taken together, these findings strongly indicate that inverted excitatory neurons exhibited an elevated intrinsic neuronal excitability, predisposing them to the integration of inputs at supra-threshold levels. Supporting this notion, our results revealed that inverted excitatory neurons exhibit a higher frequency of APs compared to upright neurons when subjected to equivalent levels of current injection.

In addition to the distinct morphological and electrophysiological features, we found that L6 upright and inverted neurons also display different synaptic properties when forming connections. Upright excitatory neurons established synaptic connections at a small inter-soma distance, whereas that of inverted neuron connections was marked larger. This may result from their different axonal projection patterns: L6 upright excitatory neurons in mPFC send their axons to subcortical areas including thalamus, claustrum, striatum, and lateral hypothalamus (Hoover and Vertes 2007). For instance, L6 corticothalamic neurons invariably show an upright dendritic morphology with an apical dendrite pointing towards the pial surface (Cotel, Fletcher et al. 2018, Hoerder-Suabedissen, Hayashi et al. 2018). However, inverted excitatory neurons are known to form intratelencephalic projections including intracortical and callosal projections (Reblet, Lopez-Medina et al. 1992, Mendizabal-Zubiaga, Reblet et al. 2007). Our axonal reconstructions showed that inverted excitatory neurons have prominent horizontal axonal projections compared to upright neurons, particularly within their home layer 6. Moreover, we found that upright excitatory neurons establish weak facilitating synapses, while synaptic connections made by inverted excitatory neurons display larger uEPSP amplitudes and show short-term depression. In rat primary somatosensory cortex, such weak and strong synapses were found to be established by corticothalamic and corticocortical excitatory neurons, respectively (Mercer, West et al. 2005, West, Mercer et al. 2006, Yang, Qi et al. 2022). These dynamic properties are specific to presynaptic neuron class, but not dependent on postsynaptic target-dependent. Our findings suggest that upright and inverted excitatory neuron populations are interconnected but functionally separable, with important implications for intracortical network function and subcortical output of mPFC layer 6.

### Adenosine modulation of the L6 microcircuitry in the mPFC

In layer 6 of the mPFC, adenosine induced a membrane potential hyperpolarisation and suppressed synaptic transmission via post- and presynaptic A_1_ARs, respectively. It has been shown that block of K_ir_ channels abolishes the adenosine-induced membrane hyperpolarisation but not the inhibition of synaptic transmission in hippocampus, brainstem and neocortex (Gerber, Greene et al. 1989, Thompson, Haas et al. 1992, Rainnie, Grunze et al. 1994, Qi, van Aerde et al. 2017). This suggests that K_ir_ channels are only involved in the postsynaptic effect of adenosine (Luscher, Jan et al. 1997, van Aerde and Feldmeyer 2015). The inhibition of synaptic transmission by adenosine via A_1_ARs is most likely due to a reduction of presynaptic neurotransmitter release resulting from a reduction in calcium influx through presynaptic voltage-dependent calcium channels (Wu and Saggau 1994, Wu and Saggau 1997). The effect of adenosine was concentration dependent and the low EC_50_ suggests that the presynaptic release probability and hence presynaptic calcium channels are more sensitive to adenosine than K_ir_ channels (Wu and Saggau 1994, van Aerde and Feldmeyer 2015, Qi, van Aerde et al. 2017).

Cell type-specific modulatory effects of adenosine via activation of A_1_ARs have been reported previously. Excitatory but not inhibitory synapses were selectively suppressed by adenosine in cortical layer 4 and in hippocampal cultures (Yoon and Rothman 1991, Qi, van Aerde et al. 2017). A_1_AR activation selectively inhibits GABA_A_ receptor currents in a specific subpopulation of hippocampal interneurons expressing axonal cannabinoid receptor type 1 (Rombo, Dias et al. 2016). However, the cell type-specific effects of adenosine on morphologically identified monosynaptic connections have not been reported yet. Applying low concentrations of adenosine (here 10 µM) to distinct excitatory synaptic connections revealed a difference in the degree of adenosine-mediated suppression between connections with a presynaptic upright and inverted neurons, respectively. Connections with an upright presynaptic neuron are more sensitive to adenosine compared to those with an inverted presynaptic neuron. This may result from a difference in the density of coupled A_1_ARs at the plasma level and/or a difference in calcium channel types at the presynaptic terminals of the two connection subtypes (Wu and Saggau 1994). However, when adenosine was applied at a concentration of 30 µM, the effect on synaptic transmission was largely similar for both types of mPFC L6 excitatory neurons. This suggests that at this higher adenosine concentration all L6 excitatory connections were strongly suppressed via A_1_AR activation, thereby masking the cell type-specific effect of adenosine.

Application of the A_1_AR antagonist CPT alone revealed the presence of endogenous extracellular adenosine in the mPFC slice preparation. In the presence of CPT alone the membrane potential became more depolarised and the spontaneous mEPSC frequency increased through a block of the tonic adenosine effect. In rat neocortex, the endogenous adenosine was estimated to be 1-2 µM, suggesting that the tonic modulation is only small and mainly affects synaptic release (Kerr, Wall et al. 2013, Qi, van Aerde et al. 2017). In hippocampus CA1 synapses, Mitchell *et al*. first provided evidence that adenosine release is activity-dependent (Mitchell, Lupica et al. 1993). A recent study in hippocampus showed that endogenous adenosine within excitatory synapses can very from 8-60 µM under basic (0.1 Hz) to high frequency (100 Hz) stimulation (Lopes, Goncalves et al. 2023). These values are much higher than the previously reported levels of extrasynaptic adenosine. An activity-dependent release of adenosine has also been found for rat neocortex (Wall and Richardson 2015). Thus, the rapid activity-dependent change in the endogenous adenosine concentration, in conjunction with its cell type- and concentration-dependent regulation of neuronal excitability and synaptic transmission reported here for layer 6 of the mPFC, indicates that adenosine plays an important role in the dynamic control of different neuronal pathways.

We also analysed the miniature spontaneous synaptic activity in L6 excitatory neurons of the mPFC. Adenosine dramatically decreased the frequency of miniature EPSCs (mEPSCs) via activation of A_1_ARs, but no changes of mEPSC amplitude was observed. Synaptic inputs onto L6 excitatory neurons originate from the thalamus or from inter- and intralaminar areas. Our finding suggests that adenosine exerts its inhibitory effect via A_1_ARs at the presynaptic terminals and reduces the neurotransmitter release probability in those cortical and subcortical areas (Fontanez and Porter 2006, Kerr, Wall et al. 2013). Previous studies on A_2A_ARs in the hippocampus have shown that activation of these receptors enhances excitatory synaptic transmission (Rebola, Pinheiro et al. 2003, Rombo, Newton et al. 2015). A study in rat visual cortex demonstrated a modulation of inhibitory transmission via A_2A_ARs, which requires the activation of A_1_ARs (Zhang, Bannon et al. 2015). However, we did not observe significant changes in either postsynaptic membrane properties or synaptic properties during application of the A_2A_AR agonist. This suggests a very low if any A_2A_ARs expression at both pre- and postsynaptic sites in layer 6 of mPFC.

Adenosine has long been implicated in the promotion and maintenance of sleep, influencing sleep spindles and slow oscillations during slow wave sleep (Landolt 2008, Bjorness, Kelly et al. 2009). Absence of compensatory slow wave sleep affected working memory function which relies on the prefrontal cortical circuits (Bjorness, Kelly et al. 2009). Sleep spindles are generated in the thalamus and are highly sensitive to modulation by thalamocortical and corticothalamic feedback circuits (Steriade 2000, Bonjean, Baker et al. 2011). Slow oscillations are generated in deep layers of the neocortex and propagate to other layers depending on recurrent intracortical synaptic activity (Massimini, Huber et al. 2004, Neske 2015). Our results show that endogenous adenosine modulates the presynaptic release of upright and inverted excitatory neurons with different sensitivity. This dynamic modulation of subcortical and intracortical synaptic transmission by adenosine may be an underlying mechanism that contributes to sleep homoeostasis. Moreover, the adenosine-induced decrease in glutamatergic synaptic transmission of L6 excitatory neurons has implications for the capacity of the mPFC to sustain persistent neural activity, a critical element supporting working memory functions (Goldman-Rakic 1995, Romo, Brody et al. 1999, Kilpatrick, Ermentrout et al. 2013). Notably, the higher adenosine sensitivity of L6 corticothalamic neurons in the PFC can affect executive functions, including planning, cognitive control, decision-making, and working memory. This influence is exerted through their projections to the mediodorsal thalamus, which serves as the primary anatomically prescribed thalamic counterpart of the PFC (Gabbott, Warner et al. 2005, Mitchell, Baxter et al. 2007, Chakraborty, Kolling et al. 2016, Bolkan, Stujenske et al. 2017, Ouhaz, Fleming et al. 2018). Further research is necessary to uncover the cell type-specific effects of adenosine on synaptic transmission. This will help to unravel the intricate functional role of adenosine in neuronal signalling in the brain.

## Author contributions

D.F., C.D. and D.Y. designed the experiments. C.D. and D.Y. carried out the patch-clamp recording experiments and electrophysiological data analysis. C.D. performed Neurolucida reconstructions and performed morphological and cluster analysis. D.Y., C.D. and D.F. wrote the manuscript. All authors have given approval for the final version of the manuscript.

## Acknowledgement

We would like to thank Werner Hucko for excellent technical assistance. We thank Dr. Karlijn van Aerde for custom-written macros in Igor Pro software. We are grateful for funding support from the European Union’s Horizon 2020 Framework Programme for Research and Innovation under the Framework Partnership Agreement No. 650003 (HBP FPA) to D.F and China Scholarship Council (to C.D.).

