## Supplementary materials for "Adenosinergic modulation of layer 6 microcircuitry in the medial prefrontal cortex is specific to presynaptic cell type"

Supplementary figures and tables

†Correspondence should be addressed to

Danqing Yang at and

Dirk Feldmeyer at

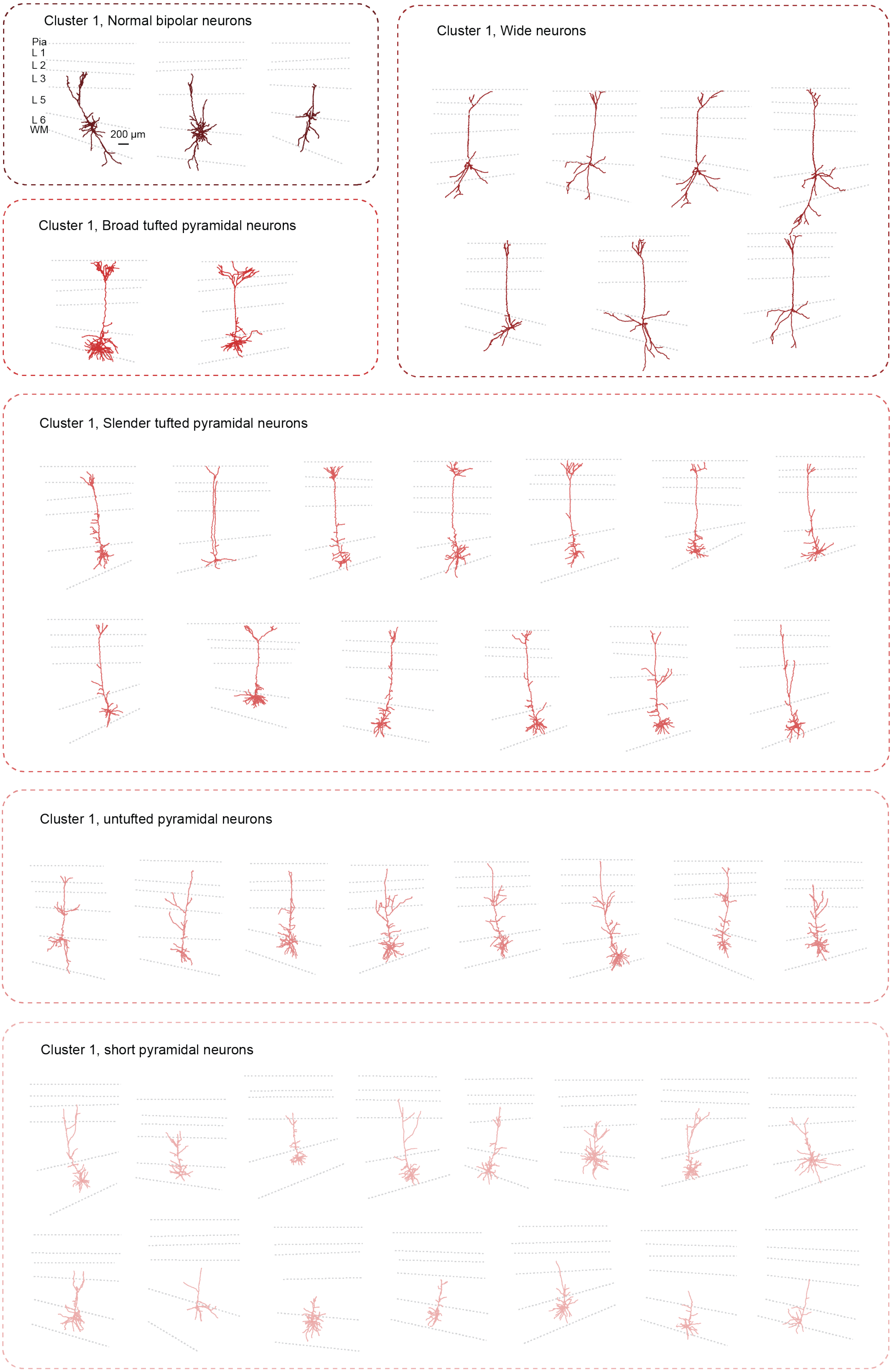

Cluster 1, Normal bipolar neurons
Pia
L 1
L 2
L 3
L 5
L 6
WM
200 µm
Cluster 1, Wide neurons
Cluster 1, Broad tufted pyramidal neurons
Cluster 1, Slender tufted pyramidal neurons
Cluster 1, untufted pyramidal neurons
Cluster 1, short pyramidal neurons

**Neurolucida reconstructions of L6 upright PNs.**

Individual morphological reconstruction ofneurons in L6 upright excitatory neuron cluster. The somatodendritic domains of distinct sub-clusters are color-coded identically as shown in **Fig. 1**. Pial surface, white matter and layer borders are indicated in dashed gray lines.

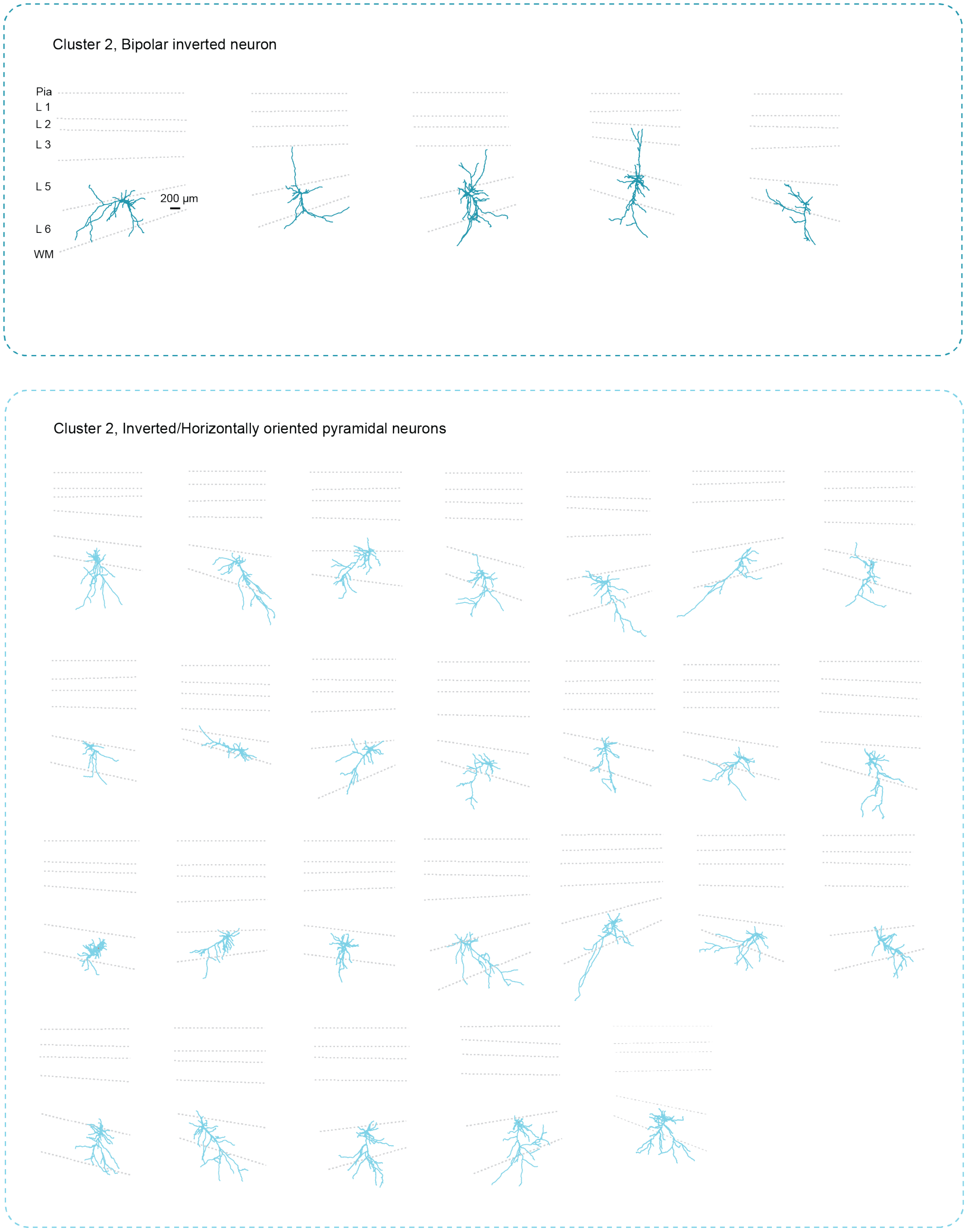


**Neurolucida reconstructions of L6 inverted neurons.**

Individual morphological reconstruction ofneurons in L6 inverted neuron cluster. The somatodendritic domains of distinct sub-clusters are color-coded identically as shown in **Fig. 1**. Pial surface, white matter and layer borders are indicated in dashed gray lines.
